# Spectinamide MBX-4888A exhibits favorable lesion and tissue distribution and promotes treatment shortening in advanced murine models of tuberculosis

**DOI:** 10.1101/2024.05.13.593953

**Authors:** Allison A. Bauman, Jansy P. Sarathy, Firat Kaya, Lisa M. Massoudi, Michael S. Scherman, Courtney Hastings, Jiuyu Liu, Min Xie, Elizabeth J. Brooks, Michelle E. Ramey, Isabelle L. Jones, Noalani D. Benedict, Madelyn R. Maclaughlin, Jake A. Miller-Dawson, Samanthi L. Waidyarachchi, Michelle M. Butler, Terry L. Bowlin, Matthew D. Zimmerman, Anne J. Lenaerts, Bernd Meibohm, Mercedes Gonzalez-Juarrero, Michael A. Lyons, Véronique Dartois, Richard E. Lee, Gregory T. Robertson

## Abstract

The spectinamides are novel, narrow-spectrum semisynthetic analogs of spectinomycin, modified to avoid intrinsic efflux by *Mycobacterium tuberculosis*. Spectinamides, including lead MBX-4888A (Lee-1810), exhibit promising therapeutic profiles in mice, as single drugs and as partner agents with other anti-tuberculosis antibiotics including rifampin and/or pyrazinamide. To demonstrate that this translates to more effective cure, we first confirmed the role of rifampin, with or without pyrazinamide, as essential to achieve effective bactericidal responses and sterilizing cure in the current standard of care regimen in chronically infected C3HeB/FeJ mice compared to BALB/c mice. Thus, demonstrating added value in testing clinically relevant regimens in murine models of increasing pathologic complexity. Next we show that MBX-4888A, given by injection with the front-line standard of care regimen, is treatment shortening in multiple murine tuberculosis infection models. The positive treatment responses to MBX-4888A combination therapy in multiple mouse models including mice exhibiting advanced pulmonary disease can be attributed to favorable distribution in tissues and lesions, retention in caseum, along with favorable effects with rifampin and pyrazinamide under conditions achieved in necrotic lesions. This study also provides an additional data point regarding the safety and tolerability of spectinamide MBX-4888A in long-term murine efficacy studies.

## Introduction

To address the ongoing global tuberculosis (TB) epidemic, there is a critical need for new shorter, more effective combination antibiotic regimens. As the cornerstone of TB control, the front-line “short-course” combination regimen consisting of an intensive treatment phase of 2-months of isoniazid (H), rifampin (R), pyrazinamide (Z) and ethambutol (E) (HRZE) and a continuation treatment phase of 4-months of HR, is deemed too lengthy to overcome problems of patient adherence, which contributes to the emergence and spread of drug-resistant strains of the etiologic agent, *Mycobacterium tuberculosis* (*Mtb*). Encouragingly, a landmark clinical trial conducted by the Tuberculosis Trials Consortium (TBTC) Study 31/AIDS Clinical Trials Group (ACTG) A5349 (Clinical_Trials.gov Identifier: NCT02410772) demonstrated non-inferiority of a four-month 4-drug regimen by replacing two drugs in the 6-month front-line regimen with the existing anti-TB agents, rifapentine and moxifloxacin (1), highlighting the potential to further shorten treatment durations for drug-susceptible *Mtb*. *In vitro* and *in vivo* preclinical models, often of increasing complexity, are used to help inform data-driven decisions on the potential of higher order TB drug combinations to achieve effective cure (2–9). One strategy successfully used to identify effective partner agents capable of contributing to treatment shortening, is to evaluate bactericidal and sterilizing activity of drug combinations in both the presence and absence of the new partner agent (for examples see, (10–13)). Regimens can then be rank ordered based on performance by one or both treatment outcomes as quantitative measures of the contribution of the new partner agent (14).

The spectinamides are a novel class of late-stage semi-synthetic antibiotics (https://www.newtbdrugs.org/pipeline/clinical) that show narrow-spectrum *in vitro* activity against both drug-susceptible and drug-resistant forms of TB (15), with promising safety and efficacy profiles *in vivo* (15–20). Key to their activity is their ability to avoid native efflux by the Rv1258c efflux pump and to selectively inhibit the mycobacterial ribosome without inhibiting mammalian mitochondrial translation (15, 21). This offers certain safety advantages over other classes of protein synthesis inhibitors, such as the oxazolidinones (22). Spectinamides have proven good partner agents for multiple TB drugs and drug regimens in a variety of murine TB efficacy models, but owing to their polar physiochemical properties, they are restricted to being used as an injectable (or inhaled) therapeutic (15, 17, 19). Particularly active with R and/or Z, spectinamides such as MBX-4888A (4888A or Lee-1810) reduced colony forming unit (CFU) burdens significantly more than R, Z or RZ alone and these *in vivo* additive effects were conserved from conventional BALB/c mice to the more advanced C3HeB/FeJ mouse model featuring advanced caseous lung pathology (17).

Here we sought to interrogate first the behavior of the drugs that make up the front-line standard of care for drug-susceptible TB in C3HeB/FeJ mice chronically infected with, relative to the conventional BALB/c model to provide context for the role of key sterilizing agents, such as R and Z, to treatment effectiveness against hard-to-treat lesion types. We also evaluated the additive effects of spectinamide MBX-4888A to the front-line regimen in terms of bactericidal response and sterilizing activity in the conventional BALB/c subacute relapsing TB infection model compared to chronically infected C3HeB/FeJ mice. Positive treatment responses to MBX-4888A combination therapy with 2HRZE/HR in multiple mouse models including mice exhibiting advanced pulmonary disease can be attributed to favorable distribution in tissues and lesions, retention in caseum, along with favorable effects with R and Z under conditions achieved in necrotic lesions *in vivo*. These encouraging drug-drug interactions between MBX-4888A and R were not recapitulated under standard *in vitro* test conditions, indicating that the effect is not purely mechanistic in nature.

## RESULTS

### Recapitulation of the “short course” regimen for TB in BALB/c and C3HeB/FeJ mice

We first confirmed the behavior of the front-line standard of care regimen in BALB/c and C3HeB/FeJ mice, chronically infected with *Mtb* Erdman, wherein the latter mouse model features advanced heterogenous lung pathology (10). The primary outcome of this study was the proportion of mice with relapse-free cure 3 months after stopping treatment. The secondary endpoint was bactericidal activity while on treatment. Lung CFU counts and proportions of mice undergoing relapse are presented in **Table 1**.

**TABLE 1.** Lung CFU counts assessed during treatment and proportion of BALB/c or C3HeB/FeJ mice relapsing after treatment completion. The numbers in brackets indicate the number of mice for which no CFU were recovered over the total number of mice at the time of plating.

| Mouse strain and treatment | Mean ± SEM lung log<sub>10</sub> CFU counts (n) | | | | | | | | | | Proportion of mice relapsing/total (%) | | | | |
| --- | --- | --- | --- | --- | --- | --- | --- | --- | --- | --- | --- | --- | --- | --- | --- |
|  | D-55 | D0 | M1 | M2 | M3 | M4 | M6 | M9 | M2 (+3) | M3 (+3) | M4 (+3) | M6 (+3) | M9 (+3) |  |  |
| **BALB/c (chronic)** |  |  |  |  |  |  |  |  |  |  |  |  |  |  |  |
| Untreated | 2.20±1.27 | 5.90 ± 0.04 (5/5) | | | | | | | | | | | | | |
| 2HRZE/HR | | | 2.62 ± 0.07 (5/5) | 0.30 ± 0.00 (3/5) | 0.30 ± 0.00 (0/5) | | | | 14/14 (100) | 14/15 (93) | | | | | |
| HE | | | | 3.84 ± 0.15 (5/5) | | | 1.18 ± 0.00 (1/5) | 1.18 ± 0.00 (0/5) | | | | 15/15 (100) | 8/15 (53) | | |
| HZE | | | | 2.69 ± 0.08 (5/5) | 1.48 ± 0.16 (5/5) | | 0.78 ± 0.00 (0/5) | | | 15/15 (100) | | 10/14 (71) | | | |
| HRE | | | | 1.21 ± 0.13 (5/5) | 0.30 ± 0.00 (2/5) | | 0.78 ± 0.00 (0/5) | | | 15/15 (100) | | 2/16 (13) | | | |
| **C3HeB/FeJ (chronic)** |  |  |  |  |  |  |  |  |  |  |  |  |  |  |  |
| Untreated | 1.51±0.81 | 7.24 ± 0.29 (5/5) | | | | | | | | | | | | | |
| 2HRZE/HR | | | 2.27 ± 0.13 (6/6) | 0.70 ± 0.26 (4/6) | 0.30 ± 0.00 (1/6) | 0.48 ± 0.00 (0/7) | | | | 12/14 (86) | 5/17 (29) | | | | |
| HE | | | | 6.35 ± 0.47 (6/6) | | | 5.51 ± 0.87 (5/6) | 5.13 ± 0.82 (5/6) | | | | 15/15 (100) | 15/15 (100) | | |
| HZE | | | | 3.93 ± 0.82 (6/6) | | 3.90 ± 1.23 (6/6) | 0.78 ± 0.00 (0/5) | | | | 12/12 (100) | 16/17 (94) | | | |
| HRE | | | | 1.81 ± 0.42 (6/6) | | 0.45 ± 0.15 (1/6) | 0.78 ± 0.00 (0/6) | | | | 12/15 (80) | 12/23 (52) | | | |

n = number of mice with CFU at plating/total D-55 = 1 day after aerosol infection; D0 = day of treatment initiation, 56 days after infection. M1 = treated for one month; M1 (+3) = treated for one month, followed by 3 months (12 weeks) without treatment, etc.

**TABLE 3.**
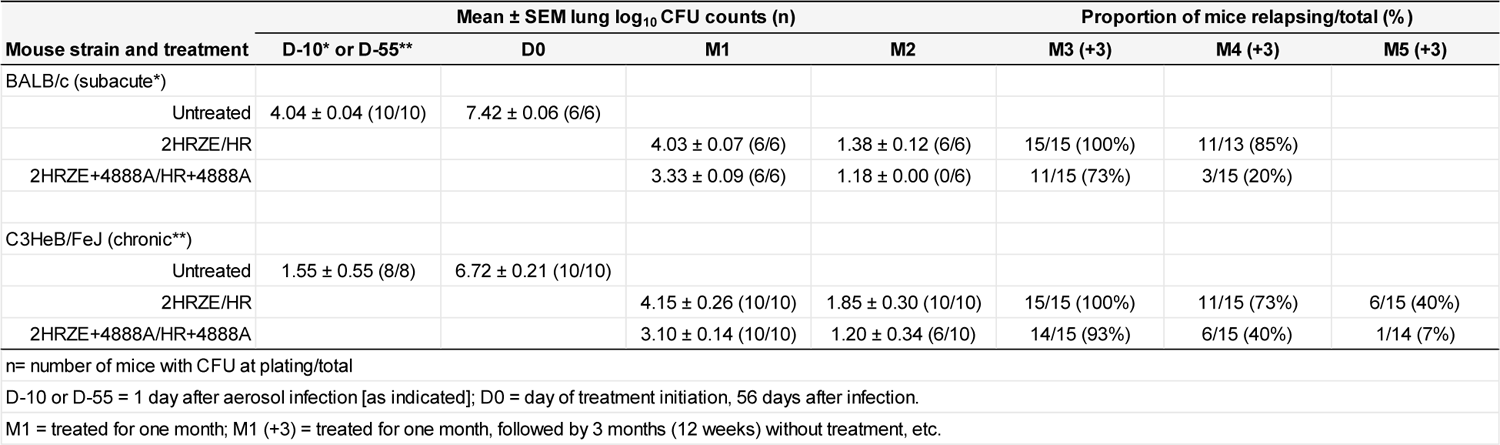
Lung CFU counts assessed during treatment and proportion of BALB/c or C3HeB/FeJ mice relapsing three months after treatment completion with the standard of care, or the standard of care augmented with spectinamide MBX-4888A.

| Mouse strain and treatment | Mean ± SEM lung $\log_{10}$ CFU counts (n) | | | | Proportion of mice relapsing/total (%) | | |
| --- | --- | --- | --- | --- | --- | --- | --- |
|  | D-10* or D-55** | D0 | M1 | M2 | M3 (+3) | M4 (+3) | M5 (+3) |
| BALB/c (subacute*) |  |  |  |  |  |  |  |
| Untreated | 4.04 ± 0.04 (10/10) | 7.42 ± 0.06 (6/6) | | | | | |
| 2HRZE/HR | | | 4.03 ± 0.07 (6/6) | 1.38 ± 0.12 (6/6) | 15/15 (100%) | 11/13 (85%) | |
| 2HRZE+4888A/HR+4888A | | | 3.33 ± 0.09 (6/6) | 1.18 ± 0.00 (0/6) | 11/15 (73%) | 3/15 (20%) | |
| C3HeB/FeJ (chronic**) |  |  |  |  |  |  |  |
| Untreated | 1.55 ± 0.55 (8/8) | 6.72 ± 0.21 (10/10) | | | | | |
| 2HRZE/HR | | | 4.15 ± 0.26 (10/10) | 1.85 ± 0.30 (10/10) | 15/15 (100%) | 11/15 (73%) | 6/15 (40%) |
| 2HRZE+4888A/HR+4888A | | | 3.10 ± 0.14 (10/10) | 1.20 ± 0.34 (6/10) | 14/15 (93%) | 6/15 (40%) | 1/14 (7%) |

n = number of mice with CFU at plating/total
D-10 or D-55 = 1 day after aerosol infection [as indicated]; D0 = day of treatment initiation, 56 days after infection.
M1 = treated for one month; M1 (+3) = treated for one month, followed by 3 months (12 weeks) without treatment, etc.

In BALB/c mice, CFU counts in lungs increased from an average log10 CFU of 2.20 (standard error of the mean [SEM] was 1.27) one day following infection to an average log10 CFU of 5.90 (SEM, 0.04) fifty-five days later at the start of treatment. As published previously for this model (10), the isoniazid-rifampin-pyrazinamide-ethambutol (HRZE) regimen reduced lung burdens by 3.28 logs with 1-month of treatment and by 5.60 logs with 2-months of treatment, wherein only 3 of 5 mice remained culture positive (**Table 1**, **Fig. 1A**). By month 3 of treatment, all mice on 2HRZE/HR were culture negative, returning no CFU within the lower limits of detection (0.3 log10 CFU per lung) (**Table 1**, **Fig. 1A**). These differences were highly significant compared to the start of treatment lung CFU controls (P < 0.0001).

**FIG 1.**
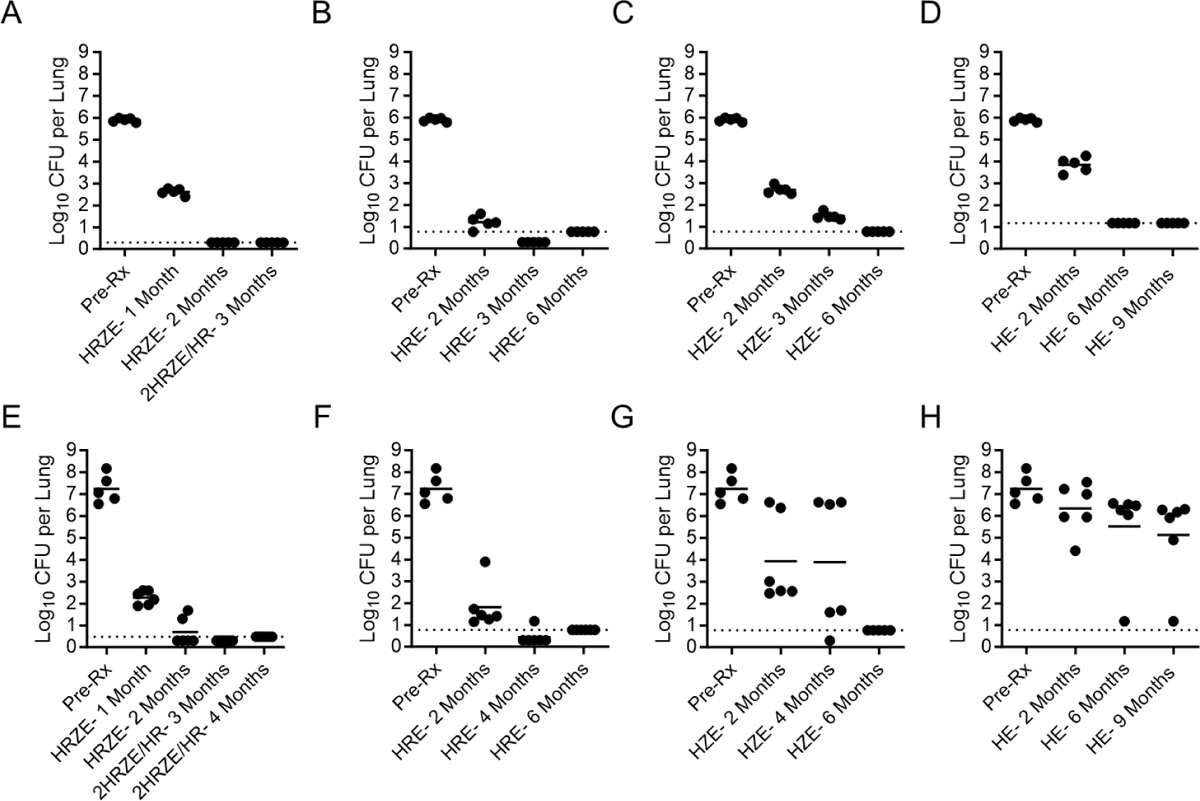
Mycobacterium tuberculosis. CFU burdens in lungs of BALB/c (A-D) and C3HeB/FeJ (E-H) mice at the start of treatment (day 0) or after treatment for the indicated number of months with 2HRZE/HR (A, E), HRE (B, F), HZE (C, G), or HE (D, H). Closed circles represent individual mice. Solid horizontal line is the group mean. Dashed horizontal line is the upper lower limit of detection.

We next evaluated the behavior of regimens lacking Z, R or both Z and R to determine the contribution of each to bactericidal responses in this model. Compared to HRZE, the HRE, HZE, and HE regimens proved significantly less bactericidal, promoting 4.69, 3.21, 2.06 log10 CFU reductions in lungs, respectively, after 2-months of treatment (**Table 1, Figs. 1B-1D**). All treatments promoted highly significant lung CFU reductions compared to the start of treatment controls (each P ≤ 0.0001). HZE and HRE continued to promote killing over time with longer treatment durations, reducing lung burdens by 4.42 and 5.60 logs after 3-months of therapy. HZE was significantly less efficacious than the standard of care, 2HRZE/HR (P<0.0001), whereas HRE could no longer be distinguished from 2HRZE/HR and only 2 of 5 mice remained culture positive with 3-months of treatment making further comparisons difficult (**Table 1, Figs. 1B-1C**). No CFU were cultured from the lungs of mice treated for 6 months with HRE or HZE, whereas 1 of 5 mice in the HE study arm remained culture positive at that timepoint. All mice were culture-negative by month 9 of HE treatment (**Table 1, Figs. 1B-1D**).

Consistent with previously published work in this model (10), chronically infected BALB/c mice treated with HRZE for 2-months or 2HRZE/HR for 3-months showed high proportions of mice relapsing after holding mice for an additional 3 months (+3) after the end of treatment (**Table 1**). All mice in the HZE and HRE treatment arms relapsed after 3-months of treatment. However, a low proportion of relapse events (2 of 16 [13%]) was observed for mice following 6-months of treatment with HRE (**Table 1**). Substitution of Z for R was significantly less sterilizing than HRE (10/14 mice relapsed [71%]; P=0.0022) (see **Table 1** and **sTable 1, Supplemental data**). HE, lacking R or Z, was the least effective regimen tested with 100% of mice relapsing following 6-months of treatment (6HE vs 6HRE, P<0.0001; 6HE vs 6HZE, P= 0.0421) (see **Table 1** and **sTable1, supplemental data**).

In C3HeB/FeJ mice, CFU counts in lungs increased from an average log10 CFU of 1.51 (SEM, 0.81) one day following infection to an average log10 CFU of 7.24 (SEM, 0.29) fifty-five days later at the start of treatment (**Table 1**). HRZE was highly effective in this model, reducing lung burdens by 4.97 logs with 1-month of treatment and by 6.54 logs with 2-months of treatment, wherein only 4 of 6 mice remained culture positive (**Table 1, Fig. 1E**). By month 3 of 2HRZE/HR treatment, only 1 of 6 mice remained culture positive (**Table 1**). These differences were highly significant compared to the start of treatment lung CFU controls (P < 0.0001) and consistent with similar effective treatment responses for 2HRZE/HR in the BALB/c arm for this same study (**Table 1**). Similar treatment responses for HRZE/HR in chronically infected C3HeB/FeJ and BALB/c mice have been reported elsewhere (10). HRE reduced C3HeB/FeJ lung burdens by 5.43 logs after 2-months of treatment (**Table 1, Fig. 1F**). Two months of treatment with HE or HZE was significantly less effective than HRZE (P<0.0001 and P=0.0018, respectively). HRE was less effective at reducing lung burdens compared to HRZE at 2-months, but the differences were not statistically significant (P = 0.4640) (**Table 1, Figs. 1E** and **1F**). As was seen in the BALB/c arm of this study, substitution of Z for R in HZE (P = 0.0482) or removing both R or Z in HE, resulted in quantitatively lower reductions in lung burdens of 3.31 and 0.89 logs, respectively, with 2-months of treatment (HE vs HRE, P < 0.0001; HE vs HZE, P =0.0214) (**Table 1, Figs. 1E-1H**). We also note a striking bi-modal treatment response in the two regimens lacking R when used to treat chronically infected C3HeB/FeJ mice, that was not apparent in BALB/c mice (compare **Figs. 1C-1D** to **Figs. 1G-1H**).

A high proportion of relapse events was observed for chronically infected C3HeB/FeJ mice after 3-months of 2HRZE/HR treatment, which was indistinguishable from the BALB/c study arm (**Table 1**). Only 5 of 17 C3HeB/FeJ mice (29%) relapsed after an additional month 2HRZE/HR (**Table 1**). These results are consistent with prior testing in this model (10). By comparison, both HZE (P = 0.0001) and HRE (P = 0.0060) were less effective than 2HRZE/HR with 100% and 80% of C3HeB/FeJ mice relapsing after 4-months of treatment (**Table 1**, and **sTable 1, supplemental data**). Substitution of Z for R in HZE resulted in a higher proportion of mice relapsing compared to the HRE arm at both 4-(not significant) and 6-months of treatment (P = 0.0053) (**Table 1**, and **sTable1, supplemental data**). Compared to the BALB/c study arm, higher proportions of C3HeB/FeJ mice relapsed following 6-months of HRE (P = 0.0173) or HZE (not significant, **Table 1**). HE was ineffective as tested here in chronically infected C3HeB/FeJ mice, with 100% of mice relapsing with 6- or 9-months of treatment (**Table 1**). The outcomes for HE appear to be mouse strain specific as HE was significantly more effective in BALB/c study arm (P = 0.0063) (**Table 1**, and **sTable 1, supplemental data**).

Co-plating indicated a limited role for expansion of drug resistance during 2HRZE/HR therapy in C3HeB/FeJ mice, with low numbers of isolates showing resistance to H, R, or E at 0.2, 1, or 5 mg/L, respectively (see **sTable 2, supplemental data**). This was similarly true for mice treated with HRE or HZE. In contrast, 5 of 6 C3HeB/FeJ mice treated with HE had high numbers of isolates capable of growth on 0.2 mg/L of H. These same mice showed a low frequency of resistance to E, suggesting the emergence of resistance to H occurred without loss of continued sensitivity to E. A higher propensity of resistance emergence was observed in the mice during relapse (see **sTable 2, supplemental data**).

### MBX-4888A contributes to bactericidal response and time to achieve durable cure in BALB/c and C3HeB/FeJ mice

Spectinamides, such as MBX-4888A, were shown previously to show strongly additive effects in combination with R and Z with 1-month of treatment *in vivo* in the BALB/c high dose aerosol subacute model and in chronically infected C3HeB/FeJ mice (17). To further explore these additive effects in murine models of increasing complexity, we next examined the contribution of adding MBX-4888A to the front-line standard of care regimen 2HRZE/HR in both the BALB/c subacute and C3HeB/FeJ chronic infection models.

The BALB/c high dose aerosol subacute TB infection model is the standard model to assess regimen performance and to enable rank-ordering of regimens for treatment shortening potential (7, 11, 12, 23–28). In this model, the bacterial load in lungs of the untreated, infected, control mice, increased from an average of 4.04 log10 CFU (SEM, 0.04) in lungs one day following aerosol infection to 7.42 log10 CFU (SEM, 0.06) in lungs at the start of treatment (Pre-Rx; 11 days post aerosol) (**Table 2**). The front-line standard of care HRZE regimen was highly effective in the BALB/c subacute TB infection model, reducing lung burdens by 3.39 and 6.04 logs with 1- or 2-months of treatment, respectively (**Table 2, Fig. 2A**). MBX-4888A given by subcutaneous injection at 200 mg/kg *quaque die* (QD), was additive with HRZE promoting a highly significant (P < 0.0001) additional 0.70 log reduction in lung burdens at 1-month, with 6 of 6 mice being culture negative with 2-months of treatment (**Table 2, Fig. 2A**). By comparison, all 6 mice in the 2-month HRZE study arm were culture positive, albeit with a low mean burden of 1.38 log10 CFU per lung (**Table 2, Fig. 2A**). Pharmacokinetic (PK) analysis of ‘in study’ sparse plasma sampling revealed no evidence of drug-drug interactions in mice receiving 4888A when it was administered alone or in combination with R and/or H (**data not shown**).

**FIG 2.**
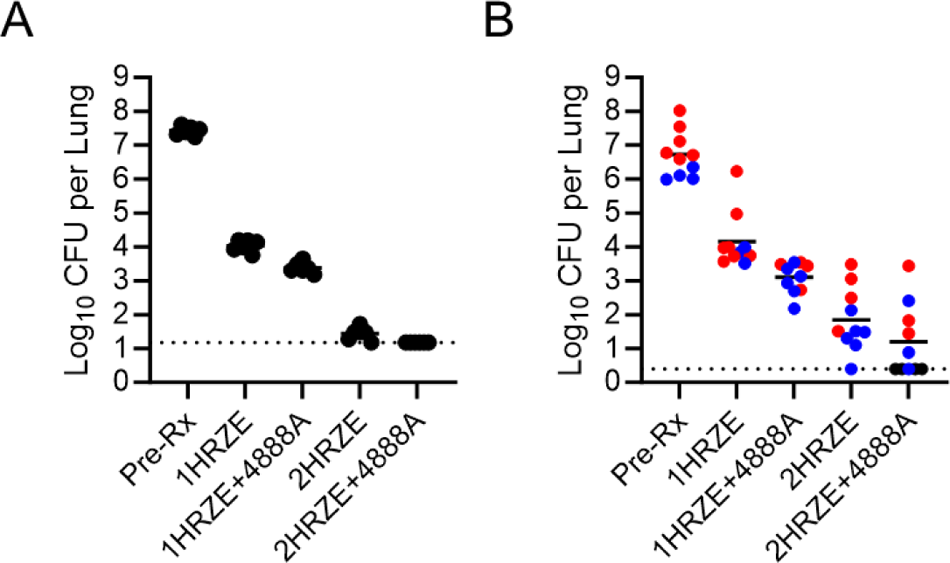
*Mycobacterium tuberculosis* CFU counts in lungs of the subacute BALB/c infection model (A) and the chronic C3HeB/FeJ (B) model at the start of treatment (day 0) or after treatment for the indicated number of months with the standard of care (2HRZE/HR) or the standard of care augmented with spectinamide MBX-4888A (2HRZE/HR+4888A). Closed circles represent individual mice. Solid horizontal line is the group mean. Dashed horizontal line is the upper lower limit of detection. Lung gross pathology based on blinded digital photographs taken at necropsy of C3HeB/FeJ mice was used to classify mice as having pronounced Type I caseous necrotic lesions (red), lacking pronounced Type I lesions and presenting with only small Type I and/or Type III inflammatory lesions (blue), or showing possible Type I lesions, which were unclear in blinded photographs (black).

The bacterial load in lungs of the untreated, infected control mice in the companion C3HeB/FeJ study arm, increased from an average of 1.55 log10 CFU (SEM, 0.55) in lungs one day following aerosol infection to 6.72 log10 CFU (SEM, 0.21) in lungs at the start of treatment (Pre-Rx; 56 days post aerosol) (**Table 2, Fig. 2**). Quantitative histopathology on 4 of 14 mice at the initiation of treatment revealed visible caseating Type I lesions in 50% of animals (see **sTable3, supplemental data**), which is in good agreement with the gross pathology assessments for the remaining mice in that study arm (**Fig. 2B**). One-month of HRZE treatment reduced the bacterial burden in the lungs of chronically infected C3HeB/FeJ mice by 2.57 log CFU, while HRZE with MBX-4888A reduced the bacterial burden in the lungs by 3.62 log CFU compared to start of treatment control mice (P < 0.0001) (see **Table 2, Fig. 2B, sTable 4, supplemental data**). Two-months of HRZE treatments also promoted significant improvements in quantitative lung pathology scores by LIRA for percent overall healthy lung tissues (78.5% vs 92.9%, P = 0.0046), and reduction in Type III lesion involvement (5.7% vs 19.5%, P = 0.0041), but had no impact, under the conditions tested herein, on the presence of Type I caseum (see **sTable 3, supplemental data**). The addition of MBX-4888A to the standard of care resulted in an improvement of 1.05 log CFU reduction in the lungs after 1-month of treatment (P=0.0026) (see **Table 2, Fig. 2B**). The addition of MBX-4888A to the standard of care did not significantly increase bactericidal responses with 2-months of treatment (P=0.1678, not significant), despite a 0.64 improvement in overall lung efficacy (HRZE vs Pre-Rx = 4.88 log CFU reduction compared to HRZE with MBX-4888A vs Pre-Rx = 5.52 log CFU reduction) (see **Table 2, Fig. 2B**). Additional net improvements including that 10 of 10 mice remained culture positive with 2-months of HRZE, while only 5 of 10 mice were culture positive when administered HRZE with MBX-4888A for 2-months are also noted (**Table 2, Fig 2B**). Thus, the addition of MBX-4888A to the standard of care resulted in an improvement of 0.64 log CFU reduction in the lungs with only half of the mice remaining culture positive after 2-months of treatment (**Table 2, Fig. 2**). From this we conclude that the addition of MBX-4888A to the standard of care regimen (HRZE) resulted in quantifiable improvements in bactericidal responses in C3HeB/FeJ mice featuring advanced lung pathology during the intensive phase of treatment.

The 2HRZE/HR regimen with and without the addition of MBX-4888A was also evaluated for sterilizing activity in C3HeB/FeJ mice with 3-, 4-, and 5-months of treatment after holding mice for an additional 3 months at the end of treatment. Five-months of treatment with the standard of care (2HRZE/HR) resulted in 40% of C3HeB/FeJ mice relapsing, with 73% and 100% of mice relapsing with 4- and 3-months of 2HRZE/HR, respectively (**Table 2**). These rates are comparable to those observed in the conventional subacute high dose aerosol BALB/c study arm and with our previous study in chronically infected BALB/c and C3HeB/FeJ mice (compare **Table 1** and **Table 2**). Addition of MBX-4888A to the standard of care resulted in a lower proportion of mice relapsing compared to 2HRZE/HR following 3-(93% compared to 100%), 4-(40% compared to 73%) or 5-months of treatment (7% compared to 40%), but these differences were not statistically significant (**Table 2** and **sTable 4, supplemental data**). Therefore, the addition of MBX-4888A to the standard of care regimen (2HRZE/HR) resulted in quantifiable improvements in achieving durable cure, that failed to reach statistical significance within the limitations of this trial design.

### Pharmacokinetics and spatial tissue distribution of MBX-4888A in C3HeB/FeJ mice

Tissue drug levels of MBX-4888A given as monotherapy by injection were next quantified by (i) standard high pressure liquid chromatography coupled to mass spectrometry (LC-MS/MS) in lung and lesion homogenates and (ii) by gravity assisted laser capture microdissection (LCM) in thin sections of specific lesion areas followed by LC-MS/MS. Four different areas were sampled by LCM within mature caseous necrotic lung granulomas: 1) lung parenchyma, lacking visible pathology, 2) the collagen rim, 3) cellular-caseum interface, consisting of foamy macrophages and neutrophils with high numbers of intracellular bacteria and 4) necrotic caseum, consisting of acellular cellular debris with high numbers of extracellular bacteria (**Fig. 3C**), shown previously to exhibit limited replication by RS ratio^®^ (28–30). MBX-4888A found drug levels were high in plasma, clearing rapidly 6-hours post dose, as published earlier (15, 18) (**Fig. 3**). Owing to favorable physiochemical properties, MBX-4888A showed excellent distribution into tissues and granuloma compartments (**Fig. 3A-3B**). MBX-4888A was further shown to be retained in tissues and lesions at levels well above the serum shifted MIC even at a time when found levels in plasma dropped below this threshold (**Fig. 3A-3B**). Similar results were obtained using both the standard LC-MS/MS approach and the highly precise LCM LC-MS/MS quantitative measure (compare **Fig. 3A** and **3B**). LCM further revealed equivalent spatial distribution across all sampled tissue compartments including caseum (**Fig. 3B**). These data confirm the hypothesis, that MBX-4888A is capable of appreciable distribution and retention into tissues including necrotic lesions, possibly reflecting favorable physiochemical attributes of the compound similar to other spectinamides (19).

**FIG. 3.**
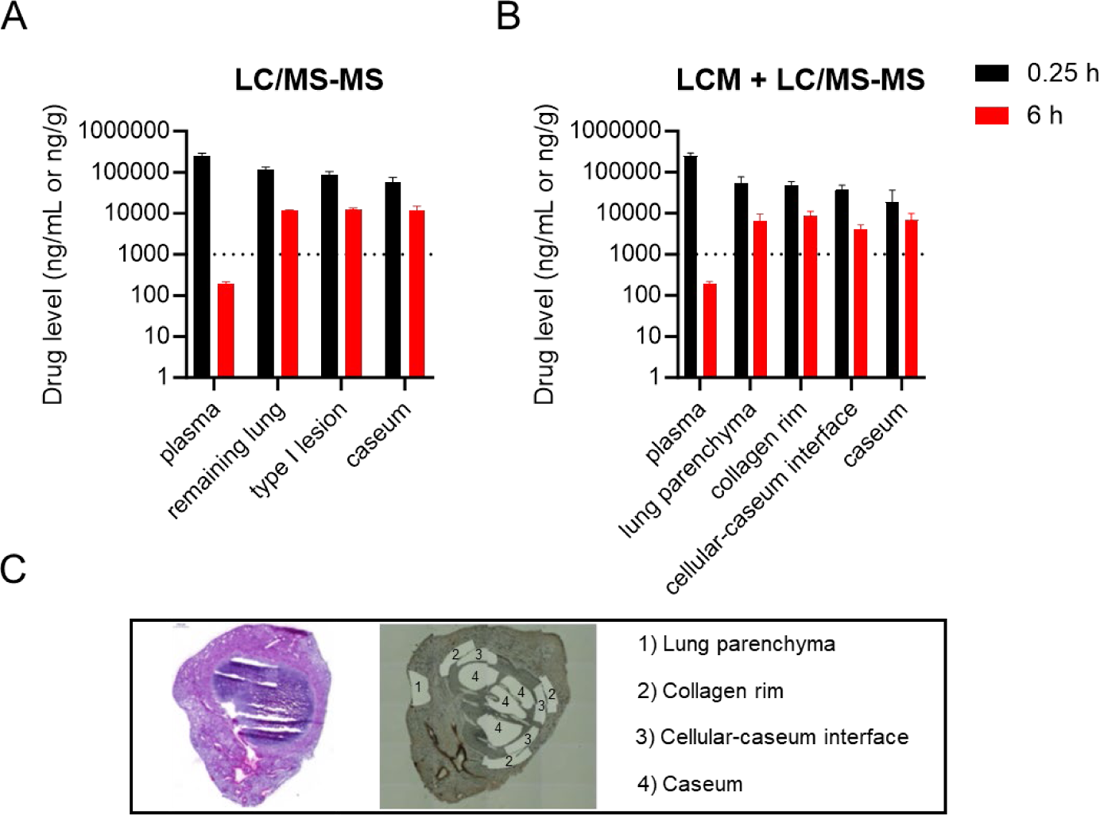
Determination of plasma and granuloma drug levels in *Mycobacterium tuberculosis* infected C3HeB/FeJ mice after 7 days of treatment with MBX-4888A (200 mg/kg, QD). Pharmacokinetic results presented are for both standard lesion LC-MS/MS (A) and LCM plus LC-MS/MS (B) approaches for quantification of spectinamide MBX-4888A in plasma and tissue samples at plasma peak (0.25 h, black) or at plasma trough time points (6 h, red). Drug levels determined by both approaches gave similar results, with high sensitivity of LCM plus LC-MS/MS enabling accurate quantification within the caseum core with a lower detection limit. The dashed line indicates the MIC in presence of 4% human serum albumin for MBX-4888A. A hematoxylin and eosin-stained lung section image is included for reference (left panel) and areas sampled by LCM are shown (right panel) (C).

### Additive effects between MBX-4888A and rifampin are not recapitulated under standard assay conditions *in vitro*

As spectinamides have been shown to exhibit mechanistic synergy *in vitro* with non-classical TB therapeutics (16), it was plausible that the MBX-4888A, through its established mechanism of action as a translational inhibitor (15), would show favorable *in vitro* interactions with R, which inhibits transcription. *In vitro* checkerboard MIC assays including a newer semi-quantitative minimum bactericidal concentration (sqMBC) endpoint (31) using *Mtb* Erdman (**Fig. 4**) or *Mtb* MC^2^ 6206 (data not shown), failed to detect any strongly favorable interactions between MBX-4888A and R, with most assays suggesting mild additivity or indifference, and no evidence of antagonism. We similarly note no interaction for MBX-4888A with H or E by these same criteria. Z was not tested owing to a lack of activity under standard assay conditions *in vitro* (32). These data are interpreted to mean that the *in vivo* additive effects described herein and elsewhere (17) are not purely mechanistic improvements in combined translation inhibition (i.e., by halting transcription and translation, simultaneously), at least not under standard checkerboard assay conditions, as was similarly reported previously (17).

**Fig. 4.**
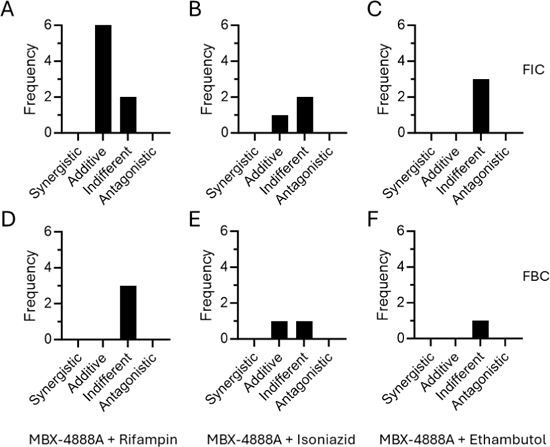
*In vitro* antimicrobial activity of MBX-4888A against *Mycobacterium tuberculosis* Erdman in broth microdilution checkerboard MIC (A-C) and semi-quantitative MBC (sqMBC) assays ((31); D-F) performed in 7H9-glycerol-ADC with 0.05% tween 80 in combination with rifampin (A, D), isoniazid (B, E), or ethambutol (C, F). Fractional inhibitory concentration (FIC) and fractional bactericidal concentration (FBC) values and frequency of occurrence showing behaviors defined herein as synergistic, FIC or FBC (≤ 0.5), additive (> 0.5 to 1), indifferent (>1 to ≤ 4), or antagonistic (>4) are shown.

### MBX-4888A is an effective partner agent against bacterial phenotypes present in caseum

The activity of MBX-4888A as a partner agent appears to be enhanced under conditions encountered in necrotic lesions (17). The central necrotic material of TB granulomas, termed caseum, is also a reservoir of drug-recalcitrant persisting mycobacteria (3, 30, 33). Previous work has suggested that *Mtb* in the caseum core exist in a slow or non-replicative phenotypic state (29, 30), which can be attributed to the hypoxic, necrotic state, due to avascularization, but also other factors such as altered pH, carbon sources, and increased reactive oxygen and nitrogen species (3, 9). These attributes contribute to drug tolerance within caseating, necrotic lesions (30), which is recapitulated in the *ex vivo* rabbit caseum MBC assay of extreme drug tolerance (33). The drug concentration kill profile of MBX-4888A, R and Z, or MBX-4888A plus R and Z was investigated in the *ex vivo* rabbit caseum MBC assay in order to isolate the ability of spectinamide MBX-4888A to contribute to killing specifically within the caseum microenvironment. Alone, MBX-4888A was weakly bactericidal in caseum with a half-maximal effective concentration (EC50) of 40 µM and dose-proportional killing with a plateau of ∼ 0.6 log10 CFU reductions at concentrations above 40 µM during monotherapy (see **Fig 5, sTables 5-6, sFig 1, supplemental data**). In contrast, R plus Z proved more effective, with EC50s of 20 µM and 400 µM, respectively and promoting dose-proportional killing exceeding 2 log10 CFU at the highest concentrations tested (see **Fig 5, sTables 5-6, sFig 2, supplemental data**). When tested in combination with R and Z, MBX-4888A enhanced this killing effect, promoting higher kills at lower drug concentrations, supporting a specific role for the spectinamide MBX-4888A, R and Z regimen combination against the bacterial phenotypes present in the caseum microenvironment (see **Fig 5, sTables 5-6, sFigs 1-3, supplemental data**).

**Fig. 5.**
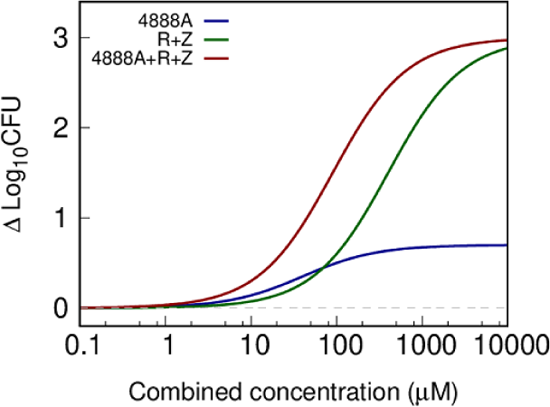
Bactericidal activity of spectinamide MBX-4888A, or rifampin (R) and pyrazinamide (Z), or MBX-4888A (4888A), rifampin and pyrazinamide (4888A+R+Z) in the *ex vivo* HN878-infected rabbit caseum assay. Data are expressed as the change from baseline in log10 of average CFU per milliliter of homogenized caseum versus combined drug concentrations from three replicates.

### MBX-4888A appears to enhance killing *in vitro* during anaerobic dormancy

Given a role for hypoxia in *Mtb* drug tolerance and phenotypic adaptations of *Mtb* in caseum due to avascularization and necrosis, we investigated whether the additive effects of spectinamide MBX-4888A with an R and Z regimen would extend to the *in vitro* rapid anaerobic dormancy model (34), which is a modification of the Wayne anaerobic dormancy model described elsewhere (35, 36). We postulated that perhaps there was a heightened need for metabolic adaptation for which the combination of these three drugs might be expected to exert a strong deleterious effect. Comparison of the biological responses 5 days after drug addition to *Mtb* adapted to dormancy under gradual oxygen depletion for 8-days in the rapid anaerobic dormancy (RAD) model, revealed expected outcomes for the positive (R) and negative controls (H), limited contribution of MBX-4888A alone, and a modest contribution of MBX-4888A when tested in combination with R or RZ for killing in the RAD model (**Fig. 6**).

**Fig. 6.**
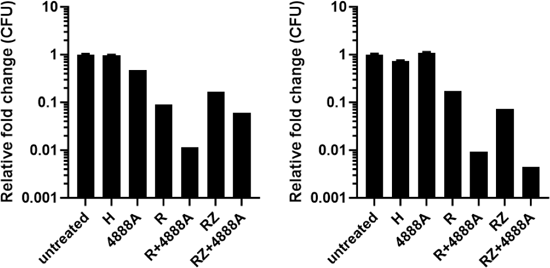
*In vitro* relative fold kill comparing the untreated (DMSO only) and drug-treated samples after 5 days of exposure to *Mycobacterium tuberculosis* adapted by gradual oxygen depletion. Isoniazid (H) was administered at 10 mg/L. Rifampin (R) and MBX-4888A (4888A) were administered at 5 mg/L. Pyrazinamide was administered at 20 mg/L. Combinations included each drug combined as described above. R and H were used as positive and negative controls, respectively, owing to their known activities in this model (37). Data are expressed as the relative fold change in CFU compared to the untreated control. Results from two independent studies with three replicates each are shown. Error bars represent the 95% CI.

## DISCUSSION

C3HeB/FeJ mice present with three distinct lesion types following virulent Mtb infection, including highly encapsulated caseous necrotic granulomas (Type I), fulminant neutrophilic alveolitis (Type II), and cellular non-necrotizing lesions (Type III), with only the latter lesion type being present in BALB/c mice (38). These models also differ in terms of the Mtb populations present, with the bacilli being predominantly extracellular in caseum, and mostly intracellular and contained within macrophages in other tissue compartments (39). The experiments presented here provide insight into the behavior of the front-line standard of care, its key drug constituents, and also improved bactericidal and sterilizing capacity when partnered with spectinamide MBX-4888A, in these two pathologically distinct murine TB efficacy models. Our central premise is that to be effective within a granulomatous lesion, drugs must be able to reach therapeutic levels for a suitable duration (i.e., PK), but importantly, they must also retain potency against the resident Mtb phenotypes arising within the different microenvironments in granulomas as well (i.e., PD) (30, 40).

Whereas, clinically effective regimens, such as 2HRZE/HR or HRE, showed nearly identical activities in BALB/c or C3HeB/FeJ mice with advanced heterogenous lung pathology, regimens lacking R or RZ (i.e., HZE or HE), were far less efficacious in C3HeB/FeJ mice, in terms of bactericidal response, prevention of relapse, and suppression of resistance emergence and outgrowth (see **Table 1** and **Figs. 1C-1D** and **Figs. 1G-1H**). Indeed, the observed rank order based on performance in both models aligns with that reported in 8-week early bactericidal activity (EBA) trials (41) and highlights the significant contribution that R or RZ adds to the current front-line regimen for drug-susceptible TB, especially in cases of hard-to-treat TB disease (42). Lanoix et al., reported a similar benefit of adding Z to HRE in terms of bactericidal response and relapse prevention in chronically infected BALB/c and C3HeB/FeJ mice (43). These findings also align with earlier results obtained in an intravenous infection model during the first two months when Z was combined with HR (44), indicating a general early benefit to adding Z to H and R containing regimens. This has also been reported in clinical studies (45).

An additional striking observation was a clear bimodal treatment response in C3HeB/FeJ mice treated with HZE or HE, that was not similarly apparent in chronically infected BALB/c mice (compare **Figs. 1C-1D** to **Figs. 1G-1H**). This phenomenon of responders and low-responders is often concordant with advanced pathology in C3HeB/FeJ mice (29, 31, 40) and is consistent with previous studies indicating R distributes at therapeutic levels and is highly active against bacterial phenotypes present in caseating lesions (30). We also note poor overall performance of HE in C3HeB/FeJ mice. The outcomes for HE appear to be mouse strain specific as HE was significantly more effective in BALB/c study arm (P = 0.0063) (**Table 1**, and **sTable1, supplemental data**). This observation is consistent with previous reports indicating limited activity of H monotherapy in C3HeB/FeJ mice (30, 39, 46) and a limited timeframe during which H benefits combination regimens in both murine and human early bactericidal assay (EBA) trials (47, 48). Conventional thinking posits that E contributes only to suppression of R-resistance in cases where the infection is resistant to H (45). Similarly, the lack of differentiation between HRZ/HR and HRZE/HR from previous work supports the generally accepted premise that E adds little to the efficacy of the HRZ-based regimen in the BALB/c relapsing mouse model (14).

Strikingly, we found that E was unable to suppress selection and outgrowth of H-resistance when administered with H in C3HeB/FeJ mice. Resistance to H emerged without concomitant selection of resistance to E, as evaluated herein. We previously reported poor performance of H in caseum during monotherapy was due to limited activity against *Mtb* phenotypes in caseum and not poor plasma to lesion distribution properties (30). We also reported appreciable selection and outgrowth of resistance to H in the C3HeB/FeJ model during monotherapy (49). However, the present data showing outgrowth of isolates exhibiting phenotypic H-resistance during HE combination therapy, does not distinguish whether this was due to periods of local monotherapy of H, or more likely, a limited ability of E to suppress and eliminate any emergent H-resistant isolates in a mouse model featuring enhanced pathology including hard-to-treat *Mtb* populations in caseum. Further studies are needed to better understand this relationship. Thus, while HE proved inferior to the other regimens tested in chronically infected BALB/c mice, as expected, the conditions encountered in C3HeB/FeJ mice chronically infected with *Mtb* revealed additional deficiencies in the ability of HE to control infection, lead to stable cure, and prevent H-resistance selection and outgrowth (see **Table 1, Fig. 1**, and **sTable 2**). These data demonstrate two strengths of the C3HeB/FeJ chronic TB mouse model, that is (i) to evaluate *in vivo* potential for resistance emergence to drugs or drug regimens in an animal model of high burden and heterogenous pathology and (ii) to identify sterilizing drugs in combinations (by reducing/tightening the bimodal responses seen by less effective combinations).

The second objective of this work was to provide additional *in vivo* efficacy data on an investigational new drug, MBX-4888A, of the spectinamide drug class. Specifically, we show here that co-administration of MBX-4888A with 2HRZE/HR improves bactericidal responses and shortens the time to achieve sufficiently low relapse events in two pathologically distinct murine TB infection models. We note that the models, as tested herein, also differ in terms of the infection status at the initiation of treatment, i.e., subacute infection [BALB/c relapsing mouse model] or chronic infection [C3HeB/FeJ model]. The present work extends a previously published study, wherein the added benefit of including a spectinamide with R or Z containing regimens was observed across three different murine TB efficacy models, including short-term C3HeB/FeJ studies (17). In contrast to that earlier work, here we also show that the confirmed early *in vivo* additive effects in terms of bactericidal response, also translates to improvements in the time it takes to achieve durable cure. In the BALB/c subacute model, adding MBX-4888A at a daily dose of 200 mg/kg QD, resulted in only 20% of mice relapsing with 4-months of treatment compared to 85% of mice administered 2HRZE/HR without MBX-4888A. Lower proportions of mice were observed to relapse when MBX-4888A was administered in addition to 2HRZE/HR in the C3HeB/FeJ mouse model featuring more human-like heterogenous pathology. Taken together, these data provide compelling evidence that spectinamide co-administration improves the bactericidal and sterilizing response rates of the front-line standard of care. This study also provides and additional data point regarding the safety and tolerability of 5 of 7 daily subcutaneous dosing of spectinamide MBX-4888A in long term murine efficacy studies. The merits and potential challenges facing co-administration of MBX-4888A, a drug that is limited at present to an injectable (15, 17, 50) or an inhaled therapeutic formulation (19, 51) with the current all oral standard of care, is beyond the scope of the current scientific study. However, long acting injectables for TB prophylaxis and treatment have recently emerged as a topic of considerable interest to the field (52).

Quantitative lung pathology scores by LIRA indicated significant improvements in lung pathology for 2HRZE/HR-treated mice compared to results obtained in companion animals at the start of treatment. Specifically, scores for overall healthy lung tissue were improved (78.5% vs 92.9%, P = 0.0046), and reductions in Type III lesion involvement (5.7% vs 19.5%, P = 0.0041) were measured, with no appreciable change on the presence of Type I caseum, which indicates limited healing of destructive, necrotic lesions, under the conditions tested herein. No mice were allocated from the 2HRZE/HR plus MBX-4888A study arm to quantitatively evaluate end of treatment pathology between these two study arms. The ability to not only quantify changes in CFU burdens, but also overall lung pathology scores during murine efficacy studies provides an additional metric by which regimen performance can be quantified that should be considered in future C3HeB/FeJ TB efficacy studies.

Drug and drug regimen effectiveness is impacted by both PK (i.e., distribution at therapeutically relevant concentrations to the site of infection) and on the *Mtb* phenotype present (i.e., drugs must exhibit activity against the *Mtb*-specific phenotype present at the site of infection) (30, 31, 53, 54). MBX-4888A showed excellent plasma to lesion distribution properties based on two complimentary analytical approaches, surgical excision and LC-MS/MS or LCM followed by LC-MS/MS. The LCM LC-MS/MS approach has the advantage of allowing highly precise drug quantification, without surgically disrupting tissues, thus avoiding possible cross-contamination between adjacent compartments. Both measures, however, provided similar results in this case, indicating favorable tissue distribution and lesion penetration properties of MBX-4888A. Indeed, MBX-4888A was retained in tissues and caseum at levels above the serum-shifted MIC far longer than in plasma (see **Fig. 3**). The excellent spatial distribution of MBX-4888A across tissues and into lesions along with retention above that of the relatively short plasma half-life of the spectinamides (15) likely contributes not only to MBX-4888A lung efficacy with QD dosing in general, but also to the overall additive effects of MBX-4888A in chronically infected C3HeB/FeJ mice featuring advanced lung pathology.

The interplay between host, pathogen, and drug PK-PD relationships, especially in complex granuloma microenvironments, is one of increasing interest to the field, and may prove impactful in data-informed regimen design to help prevent costly late-stage clinical failures (55).

We have begun to explore this dynamic in C3HeB/FeJ mice and have extended this work to include additional *ex vivo* and *in vitro* assessments that aim to simulate the *in vivo* granuloma microenvironment. In particular, the *ex vivo* caseum MBC killing assay demonstrated appreciable potency of RZ against *Mtb* in caseum, which was further enhanced by the addition MBX-4888A, which showed only limited activity on its own. In contrast, we failed to identify any obvious *in vitro* synergistic effects between MBX-4888A and R, H or E for growth inhibition or killing in conventional checkerboard assays conducted under normoxic conditions, but a general trend for improved killing with R or RZ in a rapid anaerobic dormancy model *in vitro*. Taken together, this suggests an as yet unidentified vulnerability of *Mtb* under certain conditions (i.e., anoxia or in caseum) to MBX-4888A plus R or RZ regimens, that is not purely mechanistic in nature, at least not under standard laboratory conditions. This conclusion should be considered with some degree of caution given that R was used in place of RZ, owing to the known lack of activity of Z (32) under standard *in vitro* test conditions. It is equally plausible that the effect is a combination of host-mediated effects in addition to the effects of the two drugs on the pathogen, which were not recapitulated as tested here *in vitro*. A second limitation is that while we have recently started exploration of the effect on microenvironmental stress conditions on the potency of spectinamides given as monotherapy in mice (20), we have not explored additional stress conditions shown elsewhere to promote altered drug-drug interactions and potency metrics for *in vitro* drug combination studies (5, 56). In the RAD model, cultures become sufficiently anoxic to promote conversion of methylene blue by day 6 of growth (34) as was also seen herein (data not shown). It remains unresolved whether the additive effects of MBX-4888A observed *ex vivo* in caseum, might be the environment of the caseum assay (consisting of DNA, lipids, cell debris; (3)) and not the anoxia observed in the RAD model *per se*. However, these findings highlight the potential of more advanced *ex vivo* assays such as the caseum MBC assay, currently one of our best surrogates for regimen performance in necrotic caseating granulomas *in vivo*, to help isolate drug and drug regimen performance against Mtb phenotypes found in granulomas especially when coupled to spatial drug distribution studies, as described herein.

This study has several additional limitations. First, an inherent challenge to murine relapse studies, is the need to utilize companion animals to define bactericidal versus durable cure endpoints. This limits the ability to track the overall behaviors of a given regimen to outcomes in similarly treated mice. Second, while sparse PK sampling revealed no inherent drug-drug interactions following co-administration of MBX-4888A with companion drugs tested herein, more exhaustive PK studies are needed to exclude altered drug exposures specifically related to regimen lesion distribution properties, which so far, were performed only in separate animals treated with MBX-4888A in monotherapy. Additionally, while we have focused on the known role for disease pathology altering drug distribution and efficacy, C3HeB/FeJ mice are known to exhibit increased type I interferon signaling (57), which was not considered in our study design.

In summary, this study illustrates the ability of diverse murine TB efficacy models of increasing complexity to better highlight differences in regimen behavior(s) while allowing the additive effects of individual drugs to be assessed in parallel. Specifically, we highlight inherent liabilities of removing R or RZ from the front-line TB short-course regimen in C3HeB/FeJ mice chronically infected with Mtb, in terms of reduced bactericidal response, suppression of drug-resistance, and ability to achieve durable cure, that were not apparent in the BALB/c chronic TB infection model. Such observations are highly relevant given a high incidence of resistance to R (58) and RZ (59) in human TB populations. We further demonstrate the benefit of adding a new drug, MBX-4888A, to the standard of care in both the BALB/c subacute and C3HeB/FeJ chronic TB infection models. Future studies will evaluate the contributions of each drug and pairwise combination in the HRZE+MBX-4888A regimen in a recently reported EBA-like TB murine efficacy model (called the ultra-short course model [USCM]) (60) informed based on RS ratio, an exploratory PD marker of ongoing Mtb rRNA synthesis (24, 28, 61), together with solid culture CFU counts and liquid culture time to positivity (TTP) allowing their time-course profiles to be mathematically modeled using rate equations with pharmacologically interpretable parameters. A third finding is the ability of new metrics, such as the quantitative assessment of lung pathology, and drug distribution studies to map spatial drug distribution into tissues and granulomatous lesions, to provide additional datapoints by which drug or drug regimen performance can be quantified and compared. The addition of the *ex vivo* caseum MBC assay (33), modified to accommodate drug combinations, provided additional evidence that such combinations are also capable of exerting a biological effect against the resident caseum Mtb population(s) as well. Together, such studies can be exploited to help model drug target attainment levels necessary to achieve efficacy in combination in C3HeB/FeJ mice, which holds potential to project regimen performance in human TB patients with hard-to-treat granulomatous disease.

## Materials and Methods

### Bacterial strain

*Mtb* strain Erdman (TMCC 107) was used *in vitro* and to infect mice. *Mtb* strain HN878 was used to generate rabbit caseum (33). An *Mtb* H37Rv-derived leucine/pantothenate auxotroph, MC^2^ 6206 (H37Rv Δ*leuCD* Δ*panCD*) (62), was used as a second strain for *in vitro* checkerboard assays.

### Murine Infection Models

All procedures and protocols for infecting mice with *Mtb* and subsequent drug treatments in the described mouse infection studies were approved by the Colorado State University Institutional Animal Care and Use Committee (IACUC) (Reference numbers of approved protocol: KP 1515). Mice were housed in a certified animal bio-safety level III (ABSL-3) facility. Water and mouse chow were provided *ad libitum*. Specific pathogen-free status was verified by testing sentinel mice housed within the colony. All infections were performed using a Glas-Col inhalation exposure system. In the BALB/c and C3HeB/FeJ chronic infections models, female 8- to 10-week-old mice from Jackson Laboratories (Bar Harbor, ME), were aerosol-infected using a previously calibrated *Mtb* Erdman frozen aliquot that was thawed and diluted prior to infection. The average bacterial load in lungs one-day after low-dose aerosol was approximately 2.21 log10 CFU (∼162 CFU) in BALB/c mice and ranged from 1.46 to 1.55 log10 CFU (29 to 35 CFU) for C3HeB/FeJ study arms. For the subacute TB efficacy model (high-dose aerosol model), female 6- to 8-week-old BALB/c mice (Jackson Laboratories), were aerosol-infected using *Mtb* Erdman actively grown in culture to an optical density at 600 nm (OD600) of 0.8 to 1.0 (28), resulting in the deposition of 3.8 to 4.0 log10 CFU in lungs one-day after aerosol.

In all cases, mice were block randomized and distributed equally into the different treatment arms. Infected mice were observed and weighed one- to two-times per week, due to the increased incidence of morbidity and mortality associated with clinical TB disease in the subacute BALB/c and C3HeB/FeJ mouse models. Any mice exhibiting clinical symptoms of illness were humanely euthanized.

### Antimicrobial preparation and administration to mice

Rifampin (R), isoniazid (H), and ethambutol (E) were purchased from Sigma. Pyrazinamide (Z) was purchased from Acros Organics. All four drugs were prepared in sterile water. R was prepared to achieve a final dose of 10 mg/kg after grinding to a fine powder with a mortar and pestle. H, Z, E were prepared at 1× to 3× their respective concentration needed to achieve a final dose of 10 mg/kg, 150 mg/kg, and 100 mg/kg, respectively. Z was briefly heated to 60°C until fully dissolved. MBX-4888A (4888A) was provided by Microbiotix, Inc. The 2.3 HCl salt form of 4888A was prepared in 50% (v:v) normosol:water or vitex:water solution. All drugs were prepared in weekly batches and aliquoted for daily use. All drugs were administered once daily and given 5 days (Monday through Friday) per week. R was administered by gavage in 0.2 mL volume, followed >2 h later by any companion drug (e.g., H) or drug combination. For combination studies HZ, HE, or HZE, were combined immediately before administration and given as a single oral gavage in 0.2 mL. Compound 4888A was delivered by subcutaneous injection in the flank in 0.2 mL volume at 200 mg/kg. The treatment start (D0), treatment durations (in months), and specific drug regimen employed are indicated in the study tables.

### Drug efficacy experiments and bacterial enumeration

Drug efficacy determinations were based on lung CFU counts from whole lungs or 2/3rds of the lung (by weight) aseptically harvested and frozen in 7mL Bertin Precellys tubes (CKmix50_7mL P000939-LYSK0A) 3-days after the last day of dosing to allow drug clearance from tissues. Tissues were homogenized (Precellys, Bertin Instruments, Rockville, MD) and serially diluted in PBS plus 10% bovine serum albumin. Portions of the homogenates from animals on treatment were plated for CFU on 7H11-OADC agar (i.e., Middlebrook 7H11 agar plates supplemented 0.2% [v:v] glycerol, 10% [v:v] oleic acid-albumin-dextrose-catalase (OADC) supplement, and 0.01 mg/mL cycloheximide, and 0.05 mg/mL carbenicillin) further supplemented with 0.4% [w:v] activated charcoal (7H11 charcoal agar) to help counteract drug carryover artifacts (31, 63). Lung homogenates from the relapse arms (i.e., following a 12 week [3 month] drug holiday) were plated on7H11-OADC agar without charcoal, but further supplemented with 25 mg/L polymyxin B and 20 mg/L trimethoprim to help prevent potential sample loss due to contamination. Colonies were enumerated after at least 28 days of incubation at 37°C and plates were incubated for ≥ 6 weeks to ensure all viable colonies were detected. Mice were euthanized by CO2 inhalation followed by cervical dislocation, a method approved by the IACUC at Colorado State University.

### Quantitative assessment of lung pathology at the start of treatment

Histopathology analysis was performed using a newly developed pathologist-assistive software called LIRA (‘Lesion Image Recognition and Analysis’, (64)), which reduces user limitations in terms of time and reproducibility, while providing a rapid quantitative scoring system for digital histopathology image analysis (29, 31).

### Statistical analysis

The viable CFU values were transformed to logarithms as log10 CFU per organ, which were evaluated by a one-way analysis of variance (ANOVA) with adjustments for multiple comparison using one-way Tukey’s test (pairwise comparison between all treatment groups) or Dunnett’s test (for comparison of each treatment to the start of treatment controls) using Prism 10 (GraphPad Software, San Diego, CA). Differences in relapse proportions were assessed by Fisher’s Exact test using the Holm-Bonferroni correction for multiple comparisons. Differences were considered significant at the 95% level of confidence.

### Collection of samples for pharmacokinetic analysis

For efficacy studies, plasma samples were collected during the last week of dosing to confirm the expected exposures of individual drugs when 4888A was administered with R and H.

To study 4888A PK and spatial drug distribution, C3HeB/FeJ mice were infected as above and administered 4888A starting 10-weeks post aerosol infection, to allow advanced lung pathology to fully form (see (38)). Whole blood was obtained via cardiac puncture and processed in plasma separator tubes (Becton, Dickinson and Co., Franklin Lakes, NJ) centrifuged at 3,270 RCF for 10 min at 4 °C, aliquoted into Eppendorf microcentrifuge tubes and stored at −80°C until analysis. Mice with pronounced lung pathology were used to collect samples for spatial drug quantitation by gravity assisted laser capture microdissection (LCM). Briefly, whole lung samples consisting of the cranial, medial and accessory lung lobes were collected on clear disposable base molds with the desired cutting surface in direct contact with the base of the tray (Fisher Scientific, Hampton, NH, USA). Collection trays are then placed onto a pre-chilled 4-inch aluminum block in 2 inches of liquid nitrogen in a Styrofoam cooler and allowed to sit covered for 10 minutes. Tissue trays containing frozen lobes were wrapped in foil squares, placed individually into labeled zip-lock bags and immediately transferred onto dry ice. Samples were stored at −80°C prior to analysis.

### Drug quantitation in plasma and infected tissues by HPLC coupled to tandem mass spectrometry (LC-MS/MS)

Drug levels in plasma were quantified by high pressure liquid chromatography coupled to tandem mass spectrometry (LC-MS/MS). Drug levels in tissues were measured by spatial quantitation in thin tissue sections by Laser Capture Microdissection (LCM) followed by LC-MS/MS analysis of microdissected areas (31, 65, 66).

Verapamil (IS) was purchased from Sigma Aldrich. Drug-free K2 EDTA plasma and lung tissue from CD-1 mice were obtained from BioIVT for use as blank matrices to build standard curves. A 1 mg/mL stock solution of 4888A was prepared in DMSO. Working solutions covering the desired concentration range for each drug were prepared by diluting the stock solutions in 50/50 acetonitrile (MeCN)/water. Drug-free plasma or lung homogenate (0.09 mL) were then spiked with 0.01mL of the working solutions to create the calibration standards and quality control (QC) samples. Drugs were extracted from 0.02 mL of the standard, QC and study samples by the addition of 0.2 mL of extract solvent (MeCN:MeOH, 50/50) containing 10 ng/mL verapamil. Extracts were vortexed for 5 minutes and centrifuged at 4,000 rpm for 5 minutes. An aliquot of 0.1 mL of supernatant from each specimen was transferred to a 96 well plate and 0.1 mL water was added prior to LC-MS/MS analysis.

LC-MS/MS analysis was performed on a SCIEX QTRAP 5500+ triple-quadrupole mass spectrometer coupled to a Sciex Exion UHPLC system. Chromatography was performed on an Agilent Zorbax SB-C8 column (2.1×30 mm; particle size 3.5 µm) using a reverse phase gradient. Water with 0.1% [v:v] formic acid (FA) and 0.01% [v:v] heptafluorobutyric acid (HFBA) was used for the aqueous mobile phase and 0.1% FA and 0.01% HFBA in MeCN for the organic mobile phase. Multiple-reaction monitoring (MRM) of precursor/fragment transitions in electrospray positive-ionization mode was used to quantify the analytes. MRM transitions of 469.10/263.2 and 455.40/165.00 were used for 4888A and verapamil respectively. Sample analysis was accepted if the concentrations of the quality control samples were within 20% of the nominal concentration. Data processing was performed using the Analyst software (version 1.7.2 Applied Biosystems Sciex).

### Laser-capture microdissection (LCM)

Drug levels in tissues were measured by spatial quantitation in thin tissue sections by Laser Capture Microdissection (LCM) followed by LC-MS/MS analysis of microdissected areas. Thin tissue sections were prepared and dissected as described previously (31, 65, 66). Briefly, 25 µm thick tissue sections were cut from lung biopsies using a Leica CM 1860UV cryostat (Buffalo Grove, IL) and thaw-mounted onto 1.4 µm thick Leica PET-Membrane FrameSlides (Buffalo Grove, IL). Tissue sections were immediately stored in sealed containers at −80°C. Adjacent 10 µm thick tissue sections were thaw-mounted onto standard glass microscopy slides for hematoxylin and eosin (H&E) staining.

Cellular, necrotic (caseum), and uninvolved lung lesion areas totaling 3 million µm^2^ were dissected from 1 to 3 serial lung biopsy tissue sections using a Leica LMD6 system (Buffalo Grove, IL). Areas of cellular and caseous lesion were identified optically from the brightfield image scan and by comparison to the adjacent H&E-stained section. Pooled dissected lesion tissues were collected in PCR tubes and immediately transferred to −80°C. Standard curve and QC samples were created by adding 0.01 mL of each working standard solution to 0.002 mL of 1:26.7 diluted drug-free tissue homogenates. Drugs were extracted from the calibration standard, QC, blank, and LCM study samples by the addition of 0.05 mL of extract solvent containing verapamil. Extracts were sonicated for 10 min and centrifuged at 4,000 rpm for 5 min. An aliquot of 0.05 mL of each supernatant was transferred to a 96 well plate and 0.05 mL of water was added prior to LC-MS/MS analysis.

### *In vitro* checkerboard assays

*In vitro* drug-drug interactions between 4888A and rifampin, isoniazid, or ethambutol were evaluated using a 96-well checkerboard Minimal Inhibitory Concentration (MIC) assay with a semi-quantitative Minimum Bactericidal Concentration (sqMBC) endpoint against *Mtb* Erdman in 7H9 media supplemented with 0.2% [v:v] glycerol and 10% [v:v] ADC, with 0.05% [v:v] Tween-80 (7H9 media) or *Mtb* Mc^2^ 6206 in 7H9 media further supplemented with 50 mg/L leucine, 0.2% [w:v] casamino acids, 48 mg/L pantothenate. MICs were determined by a broth microdilution assay using two-fold serial drug dilutions (31). The lowest consecutive antimicrobial concentration that showed a ≥ 80% reduction in OD600 relative to drug-free control wells, was regarded as the MIC. The sqMBC was determined on 7H11 Nunc Omni Tray (Thermo scientific, Waltham, MA) plates supplemented 0.2% [v:v] glycerol, 10% [v:v] OADC supplement (GIBCO BRL, Gaithersburg, MD), and 0.01 mg/mL cycloheximide, and 0.05 mg/mL carbenicillin with 0.4% [w:v] activated charcoal following assessment of the broth microdilution MIC on day 8 of incubation, as described previously (31).

### Caseum MBC assay

*Ex vivo* caseum bactericidal activity assay. The bactericidal activity of 4888A alone and in combination with rifampin and pyrazinamide against nonreplicating *Mtb* in rabbit caseum was conducted as described previously (33, 67). Briefly, 96-well plate wells were spotted with 4888A alone, rifampin plus pyrazinamide only, or 4888A plus rifampin and pyrazinamide to achieve the final test concentrations in **sTable 5**. Overall, each dose-response curve was designed with 4-fold increments in drug concentrations, permitting wide test ranges.

Rifampin and pyrazinamide dose ranges were centered on average concentrations achieved in the caseous compartments of lesions in patients receiving standard dosing regimens. 50 µl of 3-fold diluted caseum homogenate was added to each well in 96-well plates, which were then incubated at 37°C for 7 days. After incubation, each well was sampled and serially diluted in PBS containing 0.0125% Tween 80 prior to plating on 7H11 agar plates. Colony-forming units (CFU) were enumerated ≥ 3 weeks later.

### Drug concentration-response analysis for an *ex vivo* caseum minimum bactericidal activity assay

An Emax exposure-response model for the 4888A, R+Z, and 4888A+R+Z concentrations together with the corresponding CFU reductions, were fit to an Emax exposure-response model. The model was developed for the special case of component drug concentrations as constant ratios with a combined drug concentration, *C*, defined as

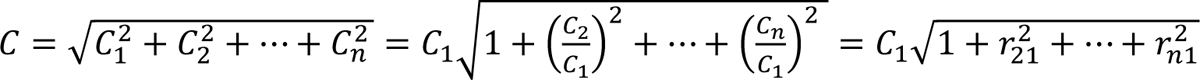

where the *C*_*i*_, *i* = 1, …, *n*, are the component drug concentrations, the *r*_*n*1_ are the ratios relative to *C*_1_, and *n* is the number of drugs in the combination. The analysis was conducted with the mean and SD CFU values. The response was calculated as reduction in log_10_(CFU) from the DMSO control as Δ log_10_(*CFU*) = log_10_(*C*FU_DMSO_) − log_10_(*C*FU), and related to the combined drug concentration, *C*, as

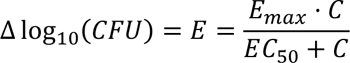

Where *E*_Max_ is the maximum effect and *EC*_50_ is the combined concentration at half-maximum effect. As the drug concentration ratios were constant, the *EC*_50,*i*_ for each drug *i* = 1, …, *n*; were calculated as

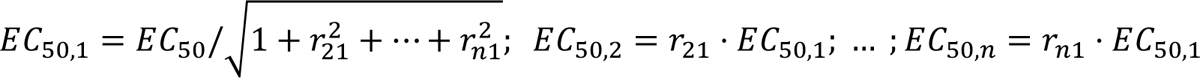

The dose-response plot included the experimental data with CFU SD error bars as

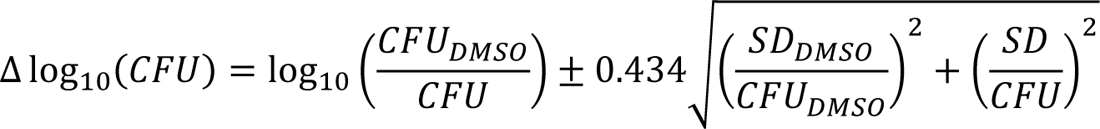

Where the 0.434 factor accounted for a change of natural to base 10 logarithm.

### Rapid Anaerobic Dormancy (RAD) assay

To test the *in vitro* activity of 4888A against *Mtb* grown under oxygen depleting conditions, the RAD assay was performed (34). Briefly, aliquots of frozen bacteria were used to start aerobic cultures in Dubos medium (Beckton-Dickinson, Sparks, MD) prepared according to manufacturer’s directions and supplemented with 0.05% Tween-80. Aerobic cultures were grown in 10 mL volumes in 150 x 25 mm screw cap tubes at 37° C with rapid stirring for 7 days, back-diluted and allowed to grow an additional 2-days to reach mid-log phase (O.D.600 = 0.4 to 0.6). Cultures were then diluted 1:100 in Dubos Tween-albumin media to an OD600 of 0.005 in 15 mm x 125 mm screw cap tubes, sealed with a butyl-rubber hungate septum stopper, and containing stir bars (12 mm by 4.5 mm), at a culture-to-headspace ratio of 0.65 (adjusted for the Colorado altitude, compared to a ratio of 0.5 for sea level). Cultures were stirred rapidly (300 rpm) using a rotary magnetic tumble stirrer (V & P Scientific, San Diego, CA). To three tubes, 0.03 mL of methylene blue stock solution at 500 mg/L was added (final concentration of 1.5 mg/L) as controls for visual confirmation of anaerobic conditions as methylene blue decolorizes with oxygen depletion (34). MBX-4888A was tested at 10×MIC (5 mg/L), as well as in combination with R (5 mg/L), Z (20 mg/L), or R and Z (5 mg/L, 20 mg/L). Isoniazid (H) was included as a negative control drug at 10 mg/L. Drugs were dissolved in 100% DMSO at 100× final concentration. Drugs were added to 8-day old anaerobic cultures (once methylene blue turned clear) by injection through the rubber septa in 0.1 mL volume. Drug exposure lasted 5 days. All compounds were tested in triplicate tubes. After 5 days of drug treatment, the tubes were uncapped and the samples suspended using a serological pipette, and serial dilutions of the cultures were plated onto 7H11 charcoal agar plates, as plating in the absence of charcoal led to drug-carryover artifacts (31, 63). Plates were incubated under normal atmospheric conditions, and bacterial colonies enumerated after ≥ 21 days of incubation at 37°C.

## Acknowledgments

This work was supported by the National Institute of Allergy and Infectious Diseases and the Office of the Director of the National Institutes of Health (grant numbers R01AI090810 [R.E.L.], R44AI098271 [Microbiotix], and 1S10OD030263 [MJG]).’ Additional support was from the Bill and Melinda Gates Foundation under grant ID numbers INV-009105, ‘TB Drug Accelerator: TB mouse *in vivo* models’ [G.T.R.], OPP1033596, “Evaluation of a New Murine Model for Testing Tuberculosis Chemotherapy” and OPP1037174, “Qualification of C3HeB/FeJ mice for Experimental Chemotherapy of Tuberculosis” [A.J.L.], and ALSAC, St. Jude Children’s Research Hospital [R.E.L]. We acknowledge Alyx M. Job and Jacy Stamps for technical contributions to this work and the staff of the Laboratory Animal Resources at Colorado State University for animal care. We are grateful to Microbiotix, Inc. (Worcester, MA), for kindly providing MBX-4888A. *Mycobacterium tuberculosis* MC^2^ 6206 was provided by William R. Jacobs, Jr. (Albert Einstein College of Medicine). The content is solely the responsibility of the authors and does not necessarily represent the official view of the National Institutes of Health.

