## supplemental data for "Spectinamide MBX-4888A exhibits favorable lesion and tissue distribution and promotes treatment shortening in advanced murine models of tuberculosis"

**Running Title:** Diverse mouse pathology impacts regimen performance

**Suppl. Table 1:** P values using the Fisher's exact test for the proportions of mice relapsing 3-months (12 weeks) after 2HRZE/HR, HRE, HZE or HE for the indicated treatment time (in months).

| BALB/c arm: Regimen/Treatment duration in months | | 2HRZE/HR | | HRE | | HZE | | HE | |
|---|---|---|---|---|---|---|---|---|---|
| | | M2+3 | M3+3 | M3+3 | M6+3 | M3+3 | M6+3 | M6+3 | M9+3 |
| 2HRZE/HR | M2+3 | NA | | | | | | | |
| | M3+3 | >0.9999 | NA | | | | | | |
| HRE | M3+3 | >0.9999 | >0.9999 | NA | | | | | |
| | M6+3 | <0.0001 | <0.0001 | <0.0001 | NA | | | | |
| HZE | M3+3 | >0.9999 | >0.9999 | >0.9999 | <0.0001 | NA | | | |
| | M6+3 | 0.0978 | 0.1686 | 0.0421 | 0.0022 | 0.0421 | NA | | |
| HE | M6+3 | >0.9999 | >0.9999 | >0.9999 | <0.0001 | >0.9999 | 0.0421 | NA | |
| | M9+3 | 0.0063 | 0.0352 | 0.0063 | 0.0233 | 0.0063 | 0.4497 | 0.0063 | NA |

| C3HeB/FeJ/c arm: Regimen/Treatment duration in months | | 2HRZE/HR | | HRE | | HZE | | HE | |
|---|---|---|---|---|---|---|---|---|---|
| | | M3+3 | M4+3 | M4+3 | M6+3 | M4+3 | M6+3 | M6+3 | M9+3 |
| 2HRZE/HR | M3+3 | NA | | | | | | | |
| | M4+3 | 0.0032 | NA | | | | | | |
| HRE | M4+3 | >0.9999 | 0.0060 | NA | | | | | |
| | M6+3 | 0.0740 | 0.2024 | 0.1006 | NA | | | | |
| HZE | M4+3 | 0.4831 | 0.0001 | 0.2308 | 0.0055 | NA | | | |
| | M6+3 | 0.5764 | 0.0002 | 0.3192 | 0.0053 | >0.9999 | NA | | |
| HE | M6+3 | 0.2241 | <0.0001 | 0.2241 | 0.0022 | >0.9999 | >0.9999 | NA | |
| | M9+3 | 0.2241 | <0.0001 | 0.2241 | 0.0022 | >0.9999 | >0.9999 | >0.9999 | NA |

**Suppl. Table 2**. Emergence of drug-resistance in the C3HeB/FeJ study arm for the indicated treatment groups at the end of each treatment arm or following a 3-month (12-week) drug-free relapse period.

| **End of treatment:** | | | | |
|---|---|---|---|---|
| Treatment Group | Timepoint | INH-R | RIF-R | EMB-R |
| 2HRZE/HR | 4 months | 0 | 0 | 0 |
| HRE | 6 months | 0 | 0 | 1 CFU (n=1) |
| HZE | 6 months | 0 | - | <0.1% (n=3) |
| HE | 9 months | 5/6* mx >UDL** (1 mx no CFU) | - | <0.1% (n=3) |

| **Relapse:** | | | | |
|---|---|---|---|---|
| Treatment Group | Timepoint | INH-R | RIF-R | EMB-R |
| 2HRZE/HR | 3mo+3obs | 3/14 (av. 10 CFU) | 12/14 (av. 500 CFU) | 3/14 (av. 1 CFU) |
| | 4mo+3obs | 0 | 0 | 0 |
| HRE | 4mo+3obs | 1/15 (av. 100 CFU) | 9/15 (av. 460 CFU) | 0 |
| | 6mo+3obs | 0 | 5/23 (av. 50 CFU) | 0 |
| HZE | 4mo+3obs | 11/12 (8mx > UDL) | - | 4/12 (av. 300 CFU) |
| | 6mo+3obs | 8/17 (>UDL) | - | 6/17 (av. 400 CFU) |
| HE | 9mo+3obs | 11/15 (8mx >UDL) | - | 1/15 (av. 1,000 CFU) |

Av., average; mx, mouse; INH-R, ≥ 0.2 mg/L; RIF-R, ≥ 1 mg/L; EMB-R, ≥ 5 mg/L

*n/N- n: mice with R-CFU, N: Total mice

**UDL- Upper Detection Limit, ~ 500 CFU/lung

**Suppl. Table 3.** Percent (%) lung involvement per lesion type at the start of treatment and after 8 2-months of HRZE treatment by LIRA.

| Mouse | Percent (%) lung involvement per lesion type at the start of treatment | | | | | | # of Type Is |
|---|---|---|---|---|---|---|---|
| | Healthy Tissue | Type I - Caseum | Type II | Type III | Type I - Rim | Unknown/Misc | |
| Mouse K | 75.7 | 3.1 | 0.0 | 17.2 | 4.0 | 0.0 | 2 |
| Mouse L | 70.8 | 0.3 | 0.0 | 28.4 | 0.5 | 0.0 | 1 |
| Mouse M | 83.8 | 0.0 | 0.0 | 16.2 | 0.0 | 0.0 | 0 |
| Mouse N | 83.2 | 0.0 | 0.0 | 16.8 | 0.0 | 0.0 | 0 |
| **Average** | **78.5** | **0.8** | **0.0** | **19.5** | **1.1** | **0.0** | **0.8** |

| Mouse | Percent (%) lung involvement per lesion type after 2-months of HRZE | | | | | | # of Type Is |
|---|---|---|---|---|---|---|---|
| | Healthy Tissue | Type I - Caseum | Type II | Type III | Type I - Rim | Unknown/Misc | |
| Mouse K | 91.4 | 0.0 | 0.0 | 8.6 | 0.0 | 0.0 | 0 |
| Mouse L | 91.5 | 2.2 | 0.0 | 4.4 | 1.9 | 0.0 | 1 |
| Mouse M | 95.9 | 0.0 | 0.0 | 4.1 | 0.0 | 0.0 | 0 |
| Mouse N | 92.7 | 0.4 | 0.0 | 5.7 | 1.2 | 0.0 | 1 |
| **Average** | **92.9** | **0.7** | **0.0** | **5.7** | **0.8** | **0.0** | **0.5** |

**Suppl. Table 4:** P values using the Fisher's exact test for the proportions of mice relapsing 3-months (12 weeks) after 2HRZE/HR or 2HRZE/HR plus MBX-4888A for the indicated treatment time (in months).

| **BALB/c arm: Regimen/Treatment duration in months** | | **HRZE/HR** | | **HRZE/HR+1810** | |
|---|---|---|---|---|---|
| | | **M3+3** | **M4+3** | **M3+3** | **M4+3** |
| **HRZE/HR** | **M3+3** | **NA** | | | |
| | **M4+3** | 0.2063 | **NA** | | |
| **HRZE/HR+4888A** | **M3+3** | 0.0996 | 0.6546 | **NA** | |
| | **M4+3** | <0.0001 | 0.0018 | 0.0092 | **NA** |

| **C3HeB/FeJ arm: Regimen/Treatment duration in months** | | **HRZE/HR** | | | **HRZE/HR+1810** | | |
|---|---|---|---|---|---|---|---|
| | | **M3+3** | **M4+3** | **M5+3** | **M3+3** | **M4+3** | **M5+3** |
| **HRZE/HR** | **M3+3** | **NA** | | | | | |
| | **M4+3** | 0.0996 | **NA** | | | | |
| | **M5+3** | 0.0007 | 0.1394 | **NA** | | | |
| **HRZE/HR+4888A** | **M3+3** | >0.9999 | 0.3295 | 0.0052 | **NA** | | |
| | **M4+3** | 0.0007 | 0.1394 | >0.9999 | 0.0052 | **NA** | |
| | **M5+3** | <0.0001 | 0.0005 | 0.0801 | <0.0001 | 0.0801 | **NA** |

**Suppl. Table 5.** Rifampin (RIF), pyrazinamide (PZA) and MBX-4888A drug concentrations evaluated in the *ex vivo* caseum bactericidal activity assay.

| | Drug concentration in µM | | |
|---|---|---|---|
| Well # | RIF | PZA | MBX-4888A |
| 1 | 0.016 | 0.313 | 0.039 |
| 2 | 0.063 | 1.250 | 0.156 |
| 3 | 0.25 | 5 | 0.625 |
| 4 | 1 | 20 | 2.5 |
| 5 | 4 | 80 | 10 |
| 6 | 16 | 320 | 40 |
| 7 | 64 | 1280 | 160 |

**Suppl. Figure 1**. Dose-response curves for spectinamide MBX-4888A in the *ex vivo* caseum bactericidal activity assay.

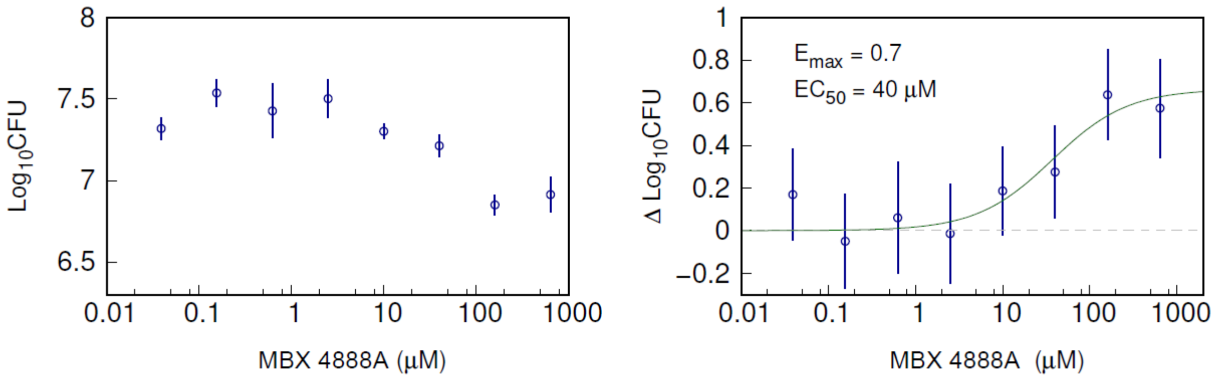


**Suppl. Figure 2**. Dose-response curves for rifampin (RIF) and pyrazinamide (PZA) in the *ex vivo* caseum bactericidal activity assay.

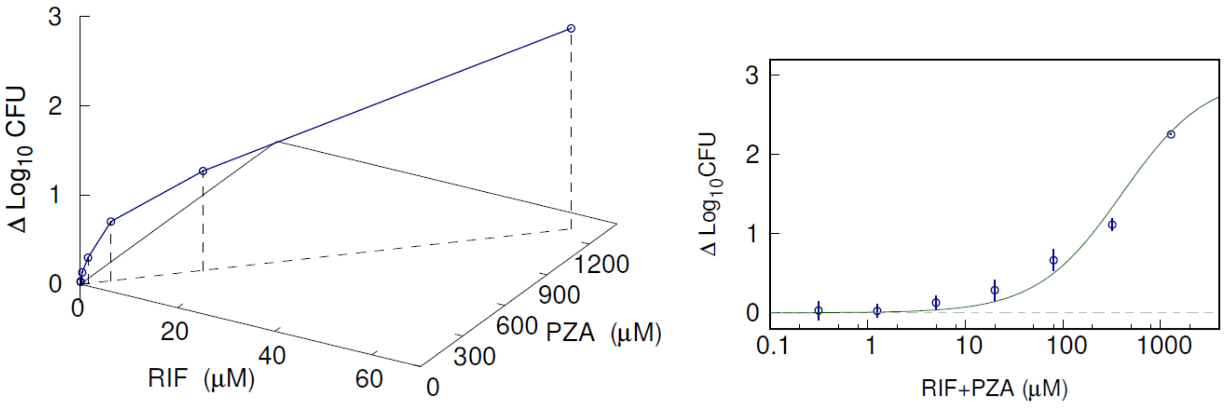

**Suppl. Figure 3**. Dose-response curves for MBX-4888A, rifampin (RIF) and pyrazinamide (PZA) in the *ex vivo* caseum bactericidal activity assay.

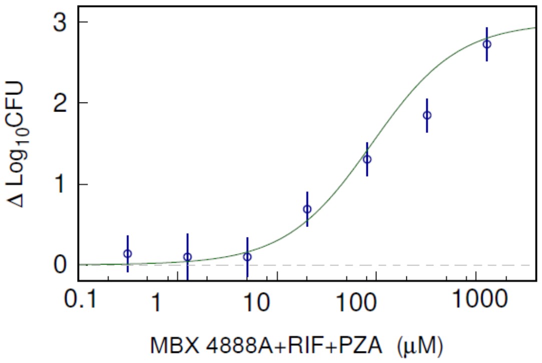


**Suppl. Table 6.** Dose-response parameters for rifampin (RIF), pyrazinamide (PZA) and MBX-4888A drug concentrations in the *ex vivo* caseum bactericidal activity assay.

| Drug | $E_{max}$ | $EC_{50}$, 4888A (μM) | $EC_{50}$, RIF (μM) | $EC_{50}$, PZA (μM) |
|---|---|---|---|---|
| MBX-4888A | 0.7 | 40 | - | - |
| RIF+PZA | 2.9 | - | 20 | 400 |
| MBX-4888A+RIF+PZA | 2.7 | 10 | 5 | 90 |
